# Embarrassingly_FASTA: Enabling Recomputable, Population-Scale Pangenomics by Reducing Commercial Genome Processing Costs from $100 to less than $1

**DOI:** 10.64898/2026.02.02.703356

**Authors:** Darren Walsh, eMalick Njie

## Abstract

Computational preprocessing has become the dominant bottleneck in genomics, frequently exceeding sequencing costs and constraining population-scale analysis, even as large repositories grow from tens of petabytes toward exabyte-scale storage to support World Genome Models. Legacy CPU-based workflows require many hours to days per 30× human genome, driving many repositories to distribute aligned or derived intermediates such as BAM and VCF files rather than raw FASTQ data. These intermediates embed reference- and model-dependent assumptions that limit reproducibility and impede reanalysis as reference genomes, including pangenomes, continue to evolve. Although recent work has established that GPUs can dramatically accelerate genomic pipelines, enabling large-cohort processing to shrink from years to days given sufficient parallelism, such workflows remain cost-prohibitive. Here, we introduce Embarrassingly_FASTA, a GPU-accelerated preprocessing pipeline built on NVIDIA Parabricks that fundamentally changes the economics of genomic data management. By rendering intermediate files transient rather than archival, Embarrassingly_FASTA enables retention of raw FASTQ data and reliable use of highly discounted ephemeral cloud infrastructure such as spot instances, reducing compute spend from ∼$17/genome (CPU on-demand) to <$1/genome (GPU spot), and commercial secondary-analysis pricing from ∼$120/genome to compute spend under $1/genome. We demonstrate the impact of this efficiency using a simulated large-cohort pangenome build-up (using variant-union accumulation as a proxy for diversity growth) in *Caenorhabditis elegans* and humans, highlighting the long tail of unsampled human genetic diversity. Beyond GPU kernels, Embarrassingly_FASTA contributes a transient-intermediate lifecycle and spot-friendly orchestration that makes FASTQ retention and routine recomputation economically viable. Embarrassingly_FASTA thus provides enabling infrastructure for recomputable, population-scale pangenomics and next-generation genomic models.

**Non-Expert Description:** Reading a person’s complete DNA sequence has become fast and inexpensive, but turning that raw data into something scientists can analyze is now one of the biggest obstacles in modern genetics. Today, processing a single genome can take many hours or even days, which makes it difficult and expensive to study large populations or reanalyze data when better methods become available. As a result, many databases store only partially processed DNA instead of the original data, limiting future discoveries.

In this work, we present a new system that dramatically speeds up this processing step using graphics processing units (GPUs), the same type of hardware used in modern artificial intelligence. With our approach, a human genome can be processed in about 35 minutes instead of more than 15 hours, and at a fraction of the cost. This makes it practical to keep the original DNA data and reprocess it whenever new tools or reference genomes become available, rather than being locked into outdated results.

We also show that this speed and affordability allow researchers to explore genetic diversity at an unprecedented scale. By analyzing both human genomes and those of a small worm species commonly used in research, we demonstrate how new genetic variations continue to emerge as more individuals are studied, especially in humans, where much diversity remains unexplored.

Overall, our work helps remove a major barrier to studying DNA at population scale and lays the foundation for future genetic models that could better explain disease, evolution, and human health.

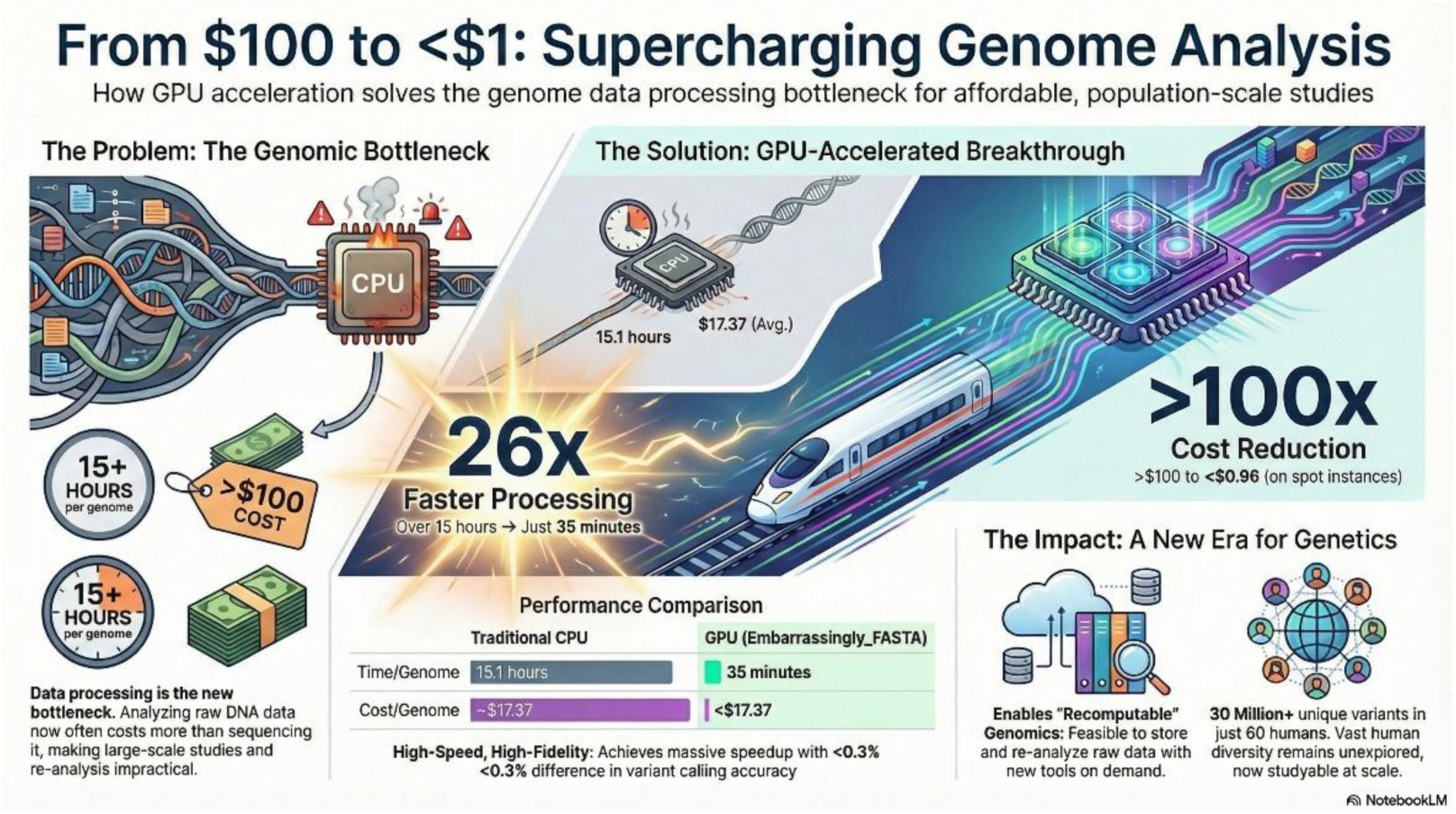

## Introduction

The cost of acquiring a human whole-genome sequence (WGS) has undergone a dramatic transformation, falling from approximately one million U.S. dollars to well below one hundred dollars today (Fig. 1). Modern sequencing platforms, however, do not output complete genomes directly. Instead, they generate unsorted collections of millions to billions of short DNA reads that must be computationally processed and assembled into a coherent genomic representation. As a result, the primary bottleneck in genomics has shifted from data generation to data processing, with computational costs now frequently exceeding sequencing costs themselves (Fig. 1).

**Figure 1:**
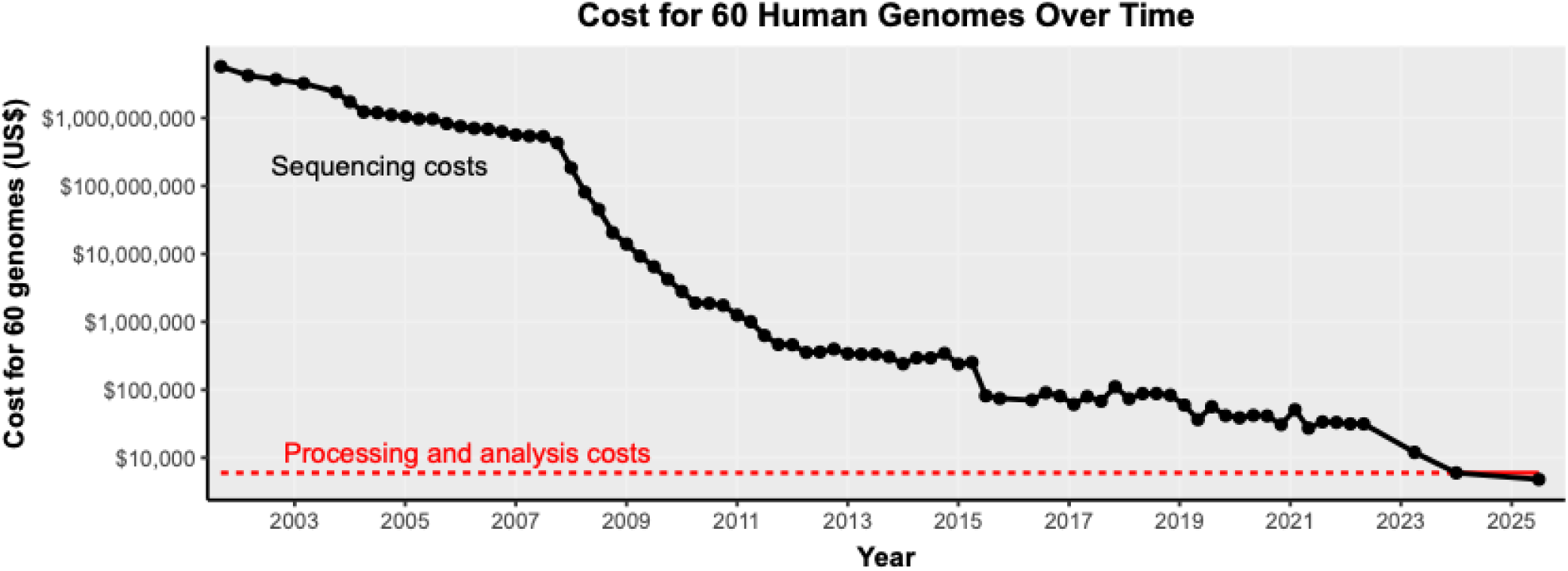
Cost of sequencing and data analysis of 60 human WGS samples over time. Sequencing costs taken from NHGRI data (Wetterstrand, 2025) and updated to include 3 latest datapoints from Illumina (*Illumina’s Revolutionary NovaSeq X Exceeds 200th Order Milestone in First Quarter 2023*, n.d.) and Ultima Genomics (*Ultima Genomics | Ultima Genomics Delivers the $100 Genome. Emerges from Stealth with $600 Million in Funding.*, 2025). Costs for data processing and analysis costs were plotted as the average ($7,260; $121 × 60 samples) of four service providers at the time of writing (January 2026).

This shift is particularly consequential for the development of World Genome Models (WGMs), which we define as population-scale genomic foundation models trained over millions of genomes to learn mechanistic representations of genetic variation across individuals, ancestries, and disease contexts (Avsec et al., 2025; Brixi et al., 2025; Dalla-Torre et al., 2025; Feng et al., 2025; Koreniuk & Njie, 2025). WGMs conceptually enable representing millions of individuals within a unified genomic substrate, such as a pangenome, while leveraging transformer-based models as a scalable computational interface for learning, generalization, and inference across populations and genomic paths. Realizing such models necessitates a fundamental expansion of genomic data infrastructure, from today’s repositories on the order of tens of petabytes toward exabyte-scale storage, sufficient to retain raw sequencing data for tens of millions of individuals and enable continual reprocessing as analytical methods evolve.

Current state-of-the-art genomic pipelines, including the Genome Analysis Toolkit (GATK), rely on CPU-intensive workflows that are prohibitive at this scale. In practice, commercial secondary analysis (FASTQ to VCF) of a single 30× human genome typically costs on the order of $120 per genome (see Methods). At population scale, this translates to billions of dollars in compute expenditure for nation-sized cohorts. Moreover, processing times of 15 hours to multiple days per genome compound to years or decades of aggregate compute, rendering routine population-scale reprocessing impractical.

These challenges are further amplified by limitations of existing reference genomes. The widely used linear reference GRCh38, as well as the more recent telomere-to-telomere assembly T2T-CHM13, both introduce reference bias by failing to represent large alternative haplotypes and by missing a substantial fraction of polymorphic structural variation present across human populations (Clavell-Revelles et al., 2025; Liao et al., 2023; Lin et al., 2024; Nyaga et al., 2025). This bias exacerbates the “street-lamp effect” in human genetics, disproportionately favoring well-represented European haplotypes while obscuring mechanisms in structurally complex or population-specific genomic regions (Clavell-Revelles et al., 2025; Liao et al., 2023; Lin et al., 2024; Nyaga et al., 2025). Graph-based pangenomes, such as those developed by the Human Pangenome Reference Consortium (HPRC), substantially improve representation of human genetic diversity by explicitly encoding alternative allelic paths (Liao et al., 2023). While the present implementation of Embarrassingly_FASTA operates over linear references, the system is explicitly designed to support graph-based pangenomic workflows as these mature, by enabling rapid recomputation across evolving reference representations. However, pangenome graphs are significantly more computationally demanding than linear references, and to date no widely adopted DNA foundation model has yet been trained directly over full graph-based human pangenomes.

To mitigate computational and storage costs, major genomic repositories, including disease-focused and clinical consortia, frequently distribute intermediate representations such as BAM and VCF files rather than the original FASTQ data generated by sequencers. While this practice reduces immediate storage and compute burdens, it introduces irreversible information loss. Alignment and variant-calling are inherently model- and reference-dependent processes that embed assumptions, heuristics, and thresholds fixed at the time of processing. Once reads are aligned and summarized, alternative alignments, multi-mapping uncertainty, complex structural signals, and non-canonical sequence features, particularly in repetitive and non-coding regions, may be permanently obscured. Although unmapped reads may be retained, the full hypothesis space of raw sequencing data is no longer recoverable, limiting reproducibility and constraining reanalysis with improved algorithms or updated reference models (Lin et al., 2024; Lloret-Villas et al., 2021).

At a computational level, the dominant operations in genomic preprocessing, sorting, aligning, and processing millions of short reads, constitute an embarrassingly parallel workload. Unlike traditional CPU pipelines, which execute these steps largely serially, Graphics Processing Units (GPUs) can exploit massive parallelism to perform thousands of independent operations concurrently. Recent studies have demonstrated order-of-magnitude speedups for read alignment and variant calling when GPU-accelerated pipelines are employed (Franke & Crowgey, 2020; O’Connell et al., 2023; Samarakoon et al., 2025).

Here, we introduce Embarrassingly_FASTA, a GPU-accelerated preprocessing pipeline built on NVIDIA Parabricks that achieves greater than 25× speedup relative to CPU-only workflows. Using 8 × NVIDIA A10 GPUs, Embarrassingly_FASTA completes end-to-end processing of a 30× human whole genome, from FASTQ to VCF, in approximately 35 minutes. This extreme throughput fundamentally alters the economics and feasibility of population-scale genomics in three ways:

i. intermediate BAM and VCF files become transient artifacts rather than archival dependencies, enabling retention of original FASTQ data without prohibitive recomputation costs;
ii. highly discounted ephemeral cloud infrastructure, such as spot instances, become reliably usable for large-scale genomic processing; and
iii. using spot instances collapses *compute spend* from ∼$17/genome (CPU on-demand) to <$1/genome (GPU spot) in our AWS configuration (compute-only; excluding storage and egress; us-east-1, Jan 2026), and reduces commercial secondary-analysis pricing (∼$120/genome) to compute spend under $1/genome.

This computational efficiency enables a new class of large-cohort analyses. Leveraging Embarrassingly_FASTA, we perform a simulated pangenome build-up to quantify how rapidly novel genetic variation is discovered as additional, non-redundant genomes are incorporated. The idealized experiment corresponds to a one-to-one mapping between samples and ancestral populations, one representative genome per distinct heritage or ecotype, such that each additional genome contributes genuinely new population-level information rather than repeated sampling of the same group. This regime is achievable in *Caenorhabditis elegans*, where curated collections provide one genome per unique isotype (ecotype), yielding a globally distributed set of distinct genomic lineages (Crombie et al., 2024). In contrast, human datasets remain necessarily coarser: even highly diverse cohorts typically consist of multiple individuals sampled from broad heritage buckets (Dalla-Torre et al., 2025; Guio et al., 2025; Jeon et al., 2024; Liao et al., 2023; Mallick et al., 2016; Suzuki, 2025), reflecting current data availability rather than true one-genome-per-heritage sampling.

Using this framework, we treat *C. elegans* as a genetically tractable proxy for a near-ideal sampling regime and humans as the best-available bucketed approximation. By comparing their variant discovery curves, we estimate how pangenome diversity scales under different sampling resolution. In this analysis, *C. elegans* shows pronounced diminishing returns by 100 ecotypes (≈3.6 million unique variant sites in this dataset), while 60 human genomes spanning five continental heritage buckets continue to add substantial novel variation with each additional genome. The contrast is consistent with differences in sampling granularity rather than a strict species-level “saturation” point. These results imply that the long tail of unsampled human heritages remains a dominant source of missing genetic variation, underscoring the necessity of truly population-scale, recomputable pangenomics.

## Results

### GPU vs CPU Performance and Variant count comparison

The GPU-accelerated pipeline achieved dramatic speedups over the legacy CPU-based workflow while maintaining comparable variant-yield metrics (Table 1).. Across eight human whole-genome sequencing (WGS) samples, the average runtime per sample on a high-end CPU server (15.1 hours) was reduced to 0.58 hours (35 minutes) on the GPU pipeline: a 26× improvement in throughput. Notably, the number of variants called by the GPU-based workflow was nearly identical to the CPU results (on the order of 5.1 million variants per genome for both). Variant counts were highly similar (<0.3% difference), indicating comparable yield. We emphasize this is a yield sanity check; rigorous correctness evaluation requires truth-set benchmarking (e.g., GIAB) or concordance analysis on shared sites. This finding is consistent with prior reports of GPU vs CPU variant concordance (99.5% for SNP calls) (O’Connell et al., 2023). In summary, each human genome was processed 25–30× faster on GPU than CPU on average, while delivering virtually the same variant yield.

**Table 1:**
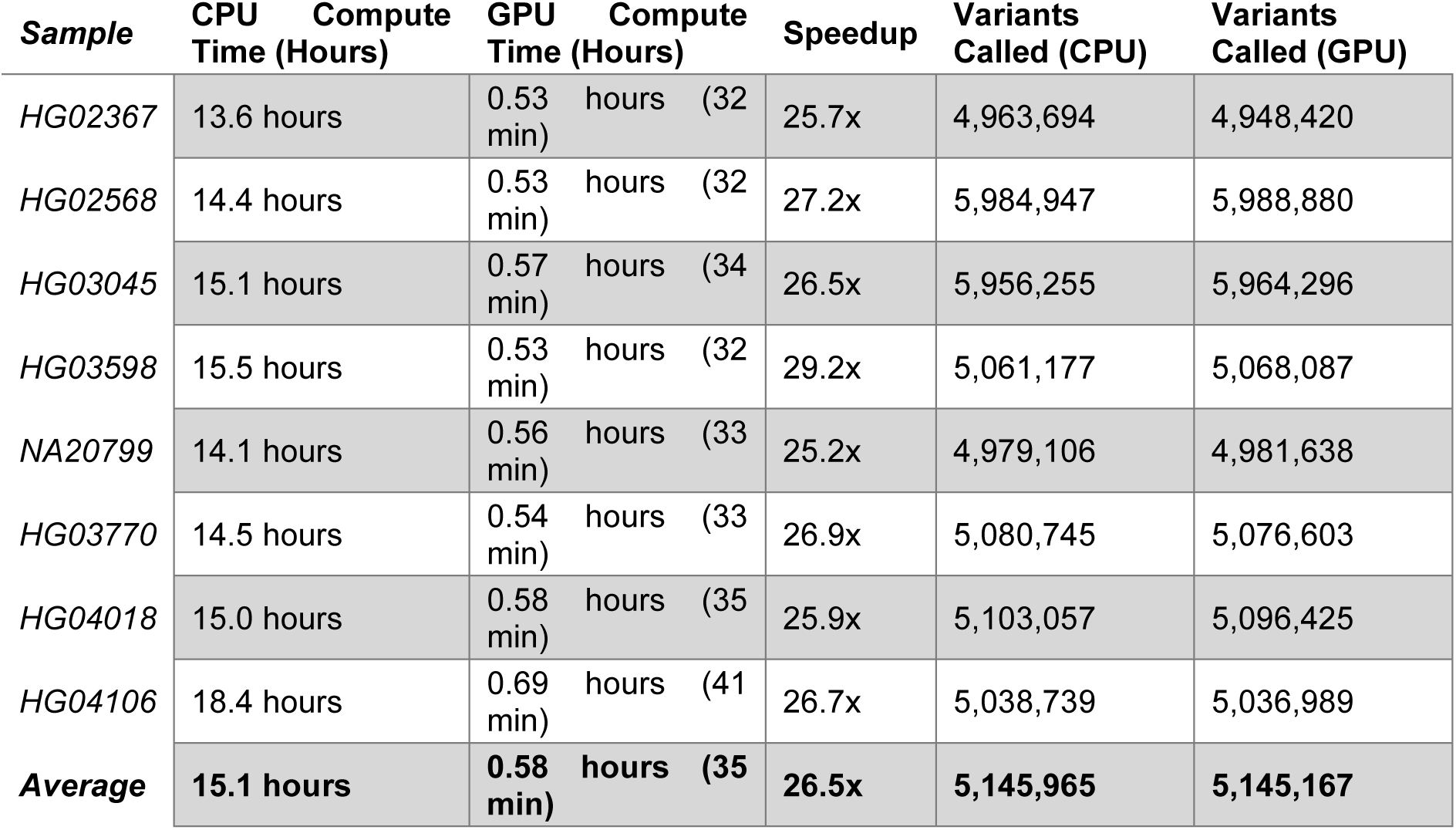
Per-sample CPU vs GPU pipeline performance for human WGS. Table comparing total compute time and variant call output for eight random representative human genomes processed with a standard CPU-based pipeline versus the GPU-accelerated pipeline. Each row is a sample (1000 Genomes Project ID); columns show CPU time (hours), GPU time (hours), the fold speedup achieved by GPU, and the number of variants called by each pipeline. The GPU runs (35 minutes per sample on average) are 25–29× faster than CPU runs (15 hours each), with an average speedup of 26.5×. Variant call yields are nearly identical between pipelines: each sample has 5 million variants, and the difference in variant counts (last two columns) is under 0.3% in all cases. These results demonstrate that GPU acceleration drastically reduces runtime while maintaining comparable variant yield (variant counts within <0.3%).

### Cross-Species Throughput (*H. sapiens* vs *C. elegans*)

The GPU workflow maintained its efficiency across different organismal genomes and scales of data (Table 2). We processed 100 *Caenorhabditis elegans* genomes (median 51 million reads each) and 60 human genomes (median 745 million reads each) to assess performance on a small versus large genome. On average, a *C. elegans* sample (100 Mb genome, 57× coverage) was analyzed in only 4.7 minutes, whereas a human sample (3.2 Gb genome, 38× coverage) took 35.8 minutes on the same GPU instance (Table 2). The larger human dataset is roughly an order of magnitude more data (mean 756 million reads vs 58 million for *C. elegans*), yet runtime increased by only 7.6×, indicating that the pipeline scales sub-linearly with input data size. This is expected as the time taken for I/O operations in *C. elegans* runs take up a larger proportion of the total processing time. The throughput remained high for human WGS at 1.7 genomes per hour per GPU instance. Average variant calls per genome reflected the species’ genomic complexity: 345,000 variants in each *C. elegans* strain versus 5.1 million in each human (Table 2). Despite these differences, the consistency of runtimes (low variance across samples) highlights robust performance of the GPU pipeline regardless of genome size or content. The genomes for *C. elegans* and humans were of ecotypes and heritages genetically and geographically distributed across the globe (Fig. 2).

**Figure 2:**
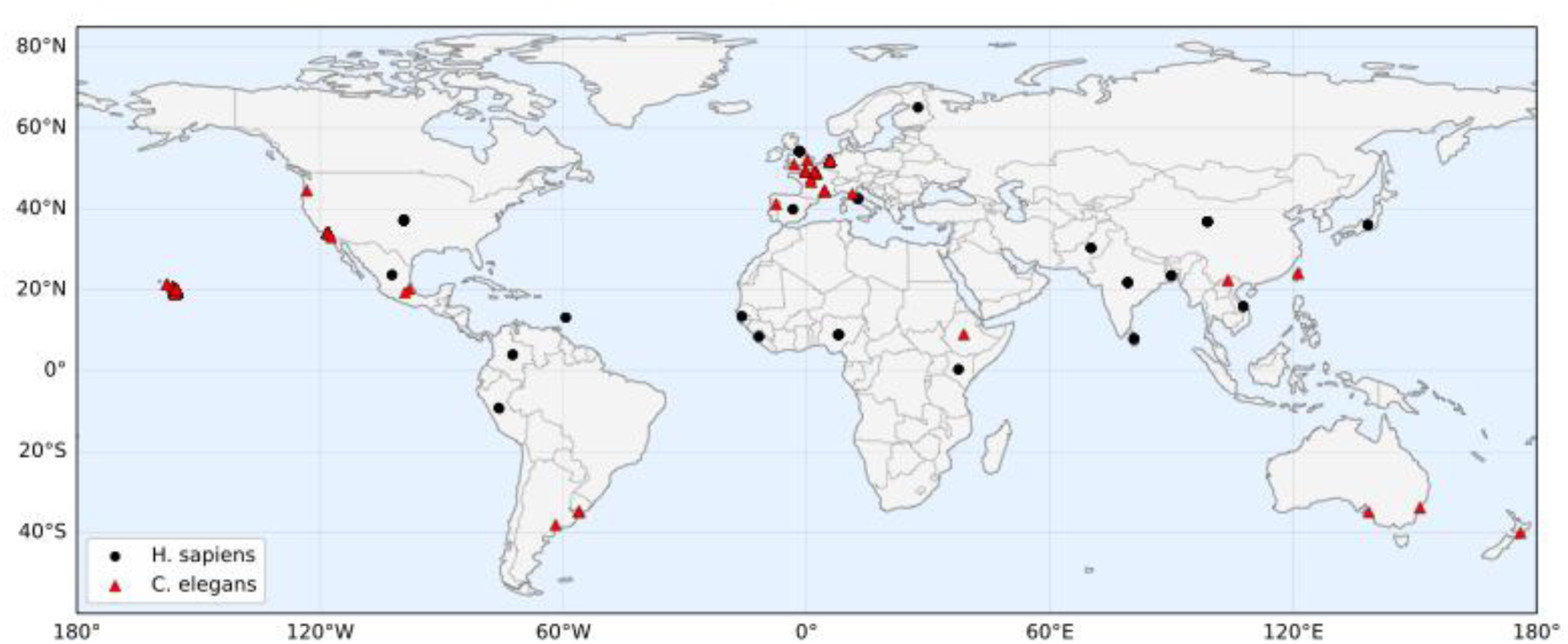
World map of sampling locations. Black circles mark *H. sapiens* locations (plotted at country centroids); red triangles mark *C. elegans* sample coordinates. Symbols match the legend in the panel. Human locations are shown at country level to respect uncertainty in individual-level provenance; *C. elegans* points reflect reported lat/long. Points with unavailable coordinates are omitted, and a minimal jitter is applied to human points when samples share the same country to reduce over-plotting. The map provides geographic context for the cohorts summarized in (a).

**Table 2:** Cross-species GPU pipeline performance and sampling geography. Aggregate metrics for *C. elegans* (n=100) and *H. sapiens* (n=60) processed with the GPU pipeline. For each species the table reports mean coverage, mean reads per sample, mean compute time per sample, and mean variants per genome. *C. elegans* (∼100 Mb genome) averaged 57× coverage, 58 M reads, 4.7 min runtime, and 3.45×10^5^ variants. *H. sapiens* (∼3.2 Gb) averaged 38× coverage, 756 M reads, 35.8 min runtime, and 5.1×10^6^ variants, illustrating consistent throughput across small and large eukaryotic genomes.

| Species | Number of Samples | Avg. Coverage | Avg. Reads Processed | Avg. Compute Time (min)<sup>1</sup> | Avg. Variants Called |
| --- | --- | --- | --- | --- | --- |
| *C. elegans* | 100 | 57x | 58 M | 4.7 min | 345,000 |
| *H. sapiens* | 60 | 38x | 756 M | 35.8 min | 5.1 M |

### Per-Sample Variability in Pipeline Performance

For the 100 *C. elegans* genomes, we further examined the distribution of input data sizes, processing times, and variant yields (Table 3). The compute times closely tracked with the number of reads per sample. For example, the fastest *C. elegans* genome run (143 seconds, 2.4 minutes) corresponded to the smallest library (16.6 million reads), while the slowest run (579 seconds, 9.7 minutes) processed the largest library (204 million reads) (Table 3). Even this worst-case scenario remained under 10 minutes. Similarly, variant counts per *C. elegans* strain ranged from 75,000 (for the least divergent strain with minimal variants) up to 749,000 (for the most variant-rich sample). The mean variant count was 345k, but the median was lower (251k), indicating a skew where a subset of strains contributed markedly more variants (perhaps wild isolates with greater divergence from the reference). Nonetheless, all 100 *C. elegans* genomes were analyzed in only a few minutes each, demonstrating that the GPU pipeline can handle large cohorts with substantial variation in coverage or variant content. The low runtimes across even the largest *C. elegans* datasets underscore the pipeline’s ability to exploit parallelism and avoid I/O bottlenecks (e.g., through on-the-fly processing), thereby efficiently scaling to hundreds of genomes. The pipeline enabled *C. elegans* samples to run 2 alignments together, thus further halving overall runtime for the full dataset.

**Table 3:**
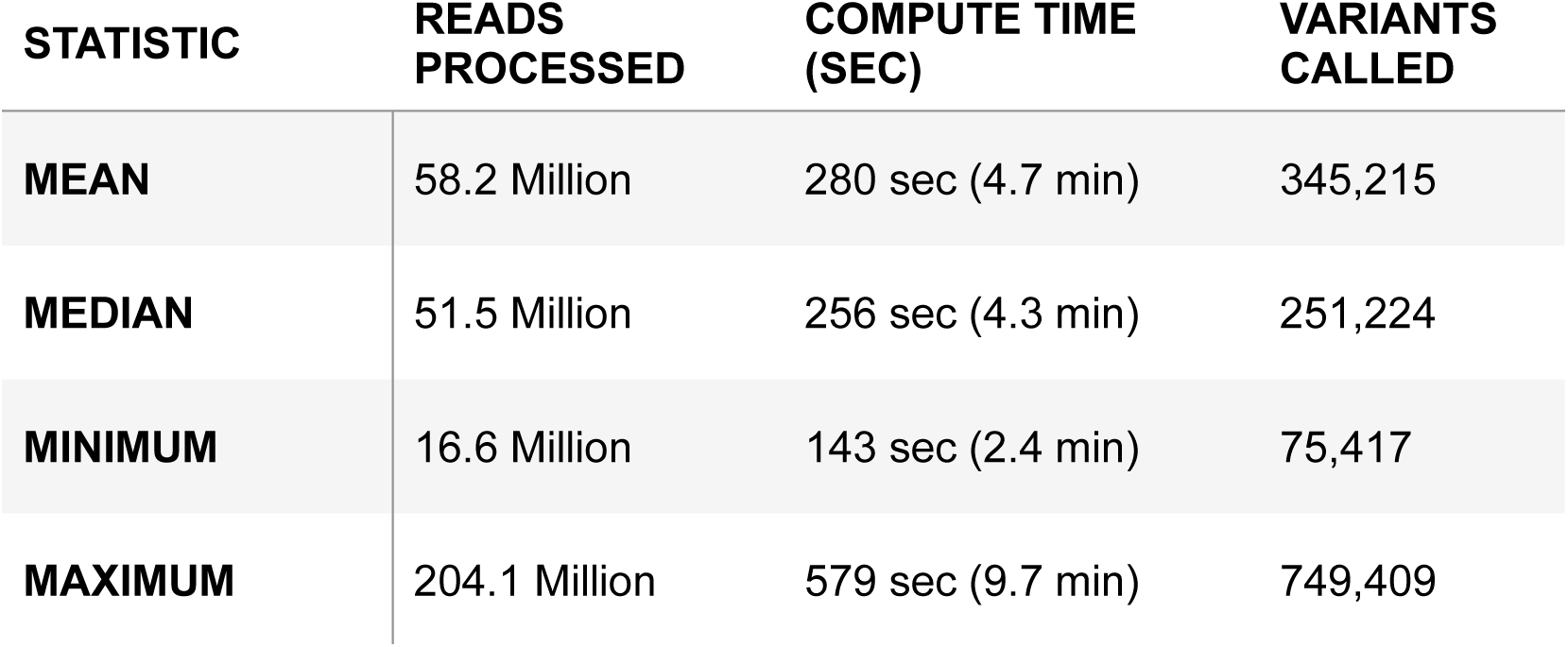
Variation in sequencing input, runtime, and variant calls across 100 *C. elegans* samples. This table provides the distribution of key metrics for the *C. elegans* cohort on the GPU pipeline: the total reads per sample, the compute time per sample (in seconds), and the number of variants called per sample. Four statistics are given: mean, median, minimum, and maximum across the 100 genomes. The mean *C. elegans* dataset was 58.2 million reads, processed in 280 seconds (4.7 min), yielding 345,215 variants. The median was slightly lower (51.5M reads, 256 sec, 251,224 variants), indicating a right-skew in coverage and variant count. The minimum sample (16.6M reads) finished in only 143 sec (2.4 min) and had 75,417 variants, whereas the largest sample (204.1M reads) took 579 sec (9.7 min) and produced 749,409 variants. These ranges show that while input data volumes and variant counts varied widely (due to biological differences and sequencing depth), even the most demanding sample was completed in under 10 minutes. The linear scaling of runtime with read count, and the proportional increase of variant calls with more reads/coverage, demonstrate the pipeline’s consistent performance and its capacity to accommodate outlier samples without bottlenecks.

### Pangenomic Diversity in *C. elegans*

We assessed how aggregate variant diversity grows as additional *C. elegans* genomes are incorporated. Fig. 3a shows the cumulative number of unique variant sites as a function of ecotypes included. The curve rises rapidly at first and then transitions to a much slower growth regime by 100 ecotypes (≈3.5–3.7 million unique variants in this dataset), indicating strong diminishing returns without a clear plateau at this cohort size. Fig. 3b shows the marginal contribution of each added ecotype: early genomes add many novel sites, but the incremental gain drops by orders of magnitude and remains non-zero at 100 genomes, consistent with continued discovery of rare and strain-specific variants with further sampling.

**Figure 3:**
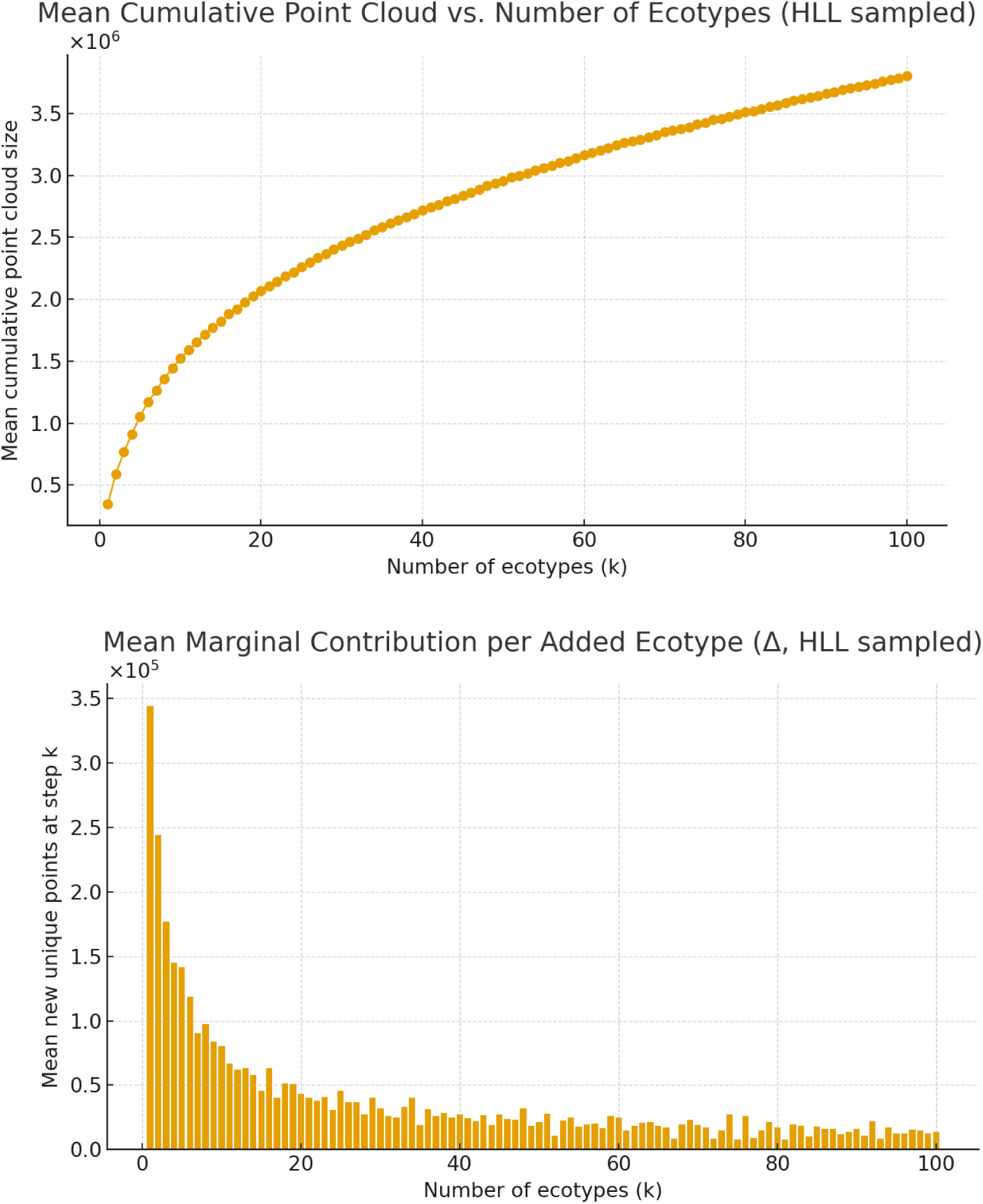
Growth of cumulative variant discovery with increasing *C. elegans* genomes (pangenome analysis). **(top)** The **mean cumulative point cloud** (total unique variant sites identified) as a function of the number of *C. elegans* ecotypes (genomes) aggregated. The curve, computed with HyperLogLog-based approximation for set cardinality, represents how the union of variant loci expands as more genomes are added. The curve rises rapidly initially and then bends toward slower growth by 100 genomes, reaching 3.6×10⁶ unique variants in this dataset. The marginal new variants per added genome decrease sharply but remain non-zero by genome 100 (on the order of 10³–10⁴), indicating ongoing accrual of rarer variants as sampling expands. This pattern is consistent with strong diminishing returns by 100 diverse strains in this dataset: many additional genomes overlap substantially with previously observed sites, while rarer and strain-specific variants continue to add new sites. **(bottom)** The mean marginal new variants per added genome (Δ unique variants) from the same data, plotted against the genome count. The first genome contributes roughly 3.5×10^5^ unique variants (points not in the reference). Each additional genome contributes fewer new variants than the previous, reflecting overlapping variation. By the 100th genome, the average marginal gain is only on the order of 10^3^ new variants. The diminishing returns indicate substantial overlap among ecotypes and progressively rarer novel sites per added genome within this cohort. Together, panels (top) and (bottom) show that cumulative diversity continues to increase with sample size, but at a much reduced rate after dozens of genomes.

Because these samples are drawn at the level of distinct ecotypes, approximating one genome per evolutionary lineage, this pronounced flattening (diminishing returns) should be interpreted primarily as an effect of sampling granularity rather than a strict upper bound on species-level diversity. Under coarser grouping (e.g., collapsing ecotypes into a small number of broad categories analogous to human continental ancestry buckets), the cumulative discovery curve would be expected to **flatten later** over the same number of samples.

Fig. 3b plots the marginal contribution of each additional genome, defined as the number of novel variant sites introduced by adding one more ecotype to the cumulative set. The first genome contributes on the order of 3.5×10⁵ unique variants, while successive genomes add progressively fewer new sites. By the time 100 ecotypes are aggregated, each additional genome contributes only a few thousand novel variants on average. However, the curve does not fully flatten, implying that variants and strain-specific loci would continue to be discovered with further sampling.

Together, these results illustrate the power of large-scale sampling in uncovering genomic diversity and foreshadow the substantially larger diversity space expected in species with greater population size and genomic complexity, such as humans. Finally, we note that all alignments were performed against a single linear reference genome, which likely suppresses the recovery of variants in regions absent or poorly represented in the reference, suggesting that true pangenomic diversity is underestimated in this analysis.

### Human WGS Performance Distribution

Table 4 provides analogous summary statistics for the 60 human genomes processed via the GPU pipeline. Each human sample had roughly 0.64–1.02 billion reads (min–max), reflecting high-coverage WGS (32× to 50×). The GPU runtimes for individual humans ranged from 1933 seconds (32.2 min) for the smallest dataset up to 2999 seconds (50.0 min) for the largest (Table 4). Despite a >1.6× difference in read count between the smallest and largest human sample, the slowest runtime was still only 50 minutes, indicating the pipeline’s strong throughput. The average runtime was 2148 sec (35.8 min), with modest variability (median 35.6 min). In terms of variant calling output, each genome yielded between 4.80 million (min) and 6.12 million (max) variants. The sample with 6.12M variants belonged to an individual from a population with higher genetic diversity (African ancestry), whereas the lower end (4.8M) corresponds to an individual with fewer divergences from the reference. The average per-genome variant count was 5.15 million (Table 4), in line with expectations for a typical human population genome at this coverage. The relatively tight distribution (median 5.03M) suggests consistent variant calling performance across samples. In summary, the GPU pipeline handled 60 human genomes in 36 minutes each on average, with runtime variation primarily driven by coverage differences, and produced variant call sets of expected size for human genetic diversity.

**Table 4:**
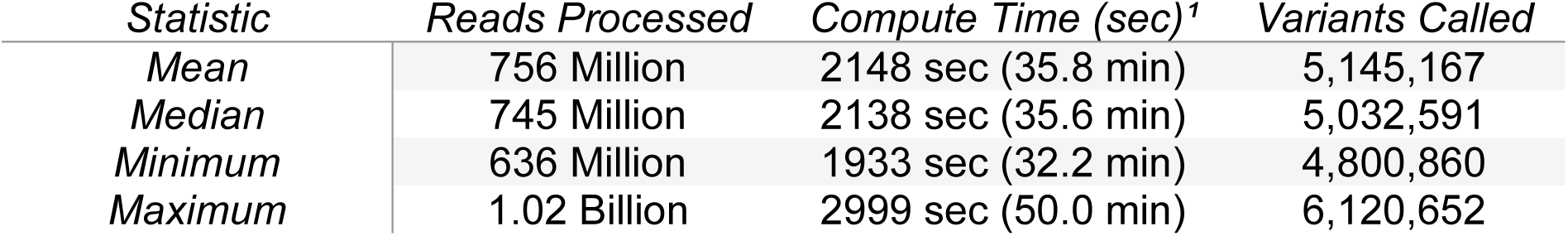
Summary of human WGS input size, runtime, and variant count distributions on GPU. This table mirrors Figure 3 but for 60 human genomes processed by the GPU pipeline. It reports the mean, median, minimum, and maximum per-sample values for total reads, compute time, and variants called. On average, each human genome had 756 million reads (38× coverage), took 2148 seconds (35.8 min) to analyze, and yielded 5,145,167 variant calls. The median was similar: 745M reads, 2138 sec (35.6 min), and 5,032,591 variants. The fastest run (minimum time 1933 sec, 32.2 min) corresponded to the smallest dataset (636M reads) and produced 4.80M variants. The slowest run (max time 2999 sec, 50.0 min) was for the largest dataset (1.02 billion reads) with 6.12M variants called. The relatively narrow spread between mean and median indicates most samples clustered around the 35–36 minute mark and 5 million variants. Outlier genomes with higher coverage proportionally took longer and yielded more variants, but notably even 50 minutes is a modest runtime for a 1 billion-read genome. This figure underscores the consistent, scalable performance of the GPU pipeline for human WGS, and that per-genome variant output falls in expected ranges (approximately 4.8–6.1 million) depending on individual genetic diversity.

### Cumulative Human Genetic Diversity

We next examined how the union of variant sites grows as multiple human genomes are aggregated. Fig. 4a shows the mean cumulative variant count as a function of the number of human genomes combined (from 1 up to 60), using an approximate cardinality estimator to count unique sites. Starting from a single genome (5 million variants), the total unique variant count increased steeply with each additional genome and reached 30 million after combining all 60 individuals. Notably, the curve has not reached a clear plateau at 60; it continues to climb, indicating that substantial novel variation is introduced even by the later genomes. This reflects the fact that human populations harbour enormous diversity and even 60 genomes are far from capturing all common variants, let alone rare variants. Fig. 4b highlights the diminishing but non-zero returns: the first genome contributes 5×10^6^ unique variant sites (by definition), the second adds roughly 2–3 million new sites (not present in the first), and the incremental gain per genome keeps decreasing thereafter. By the 60th genome, the marginal new variants contributed are on the order of only tens of thousands.

**Figure 4:**
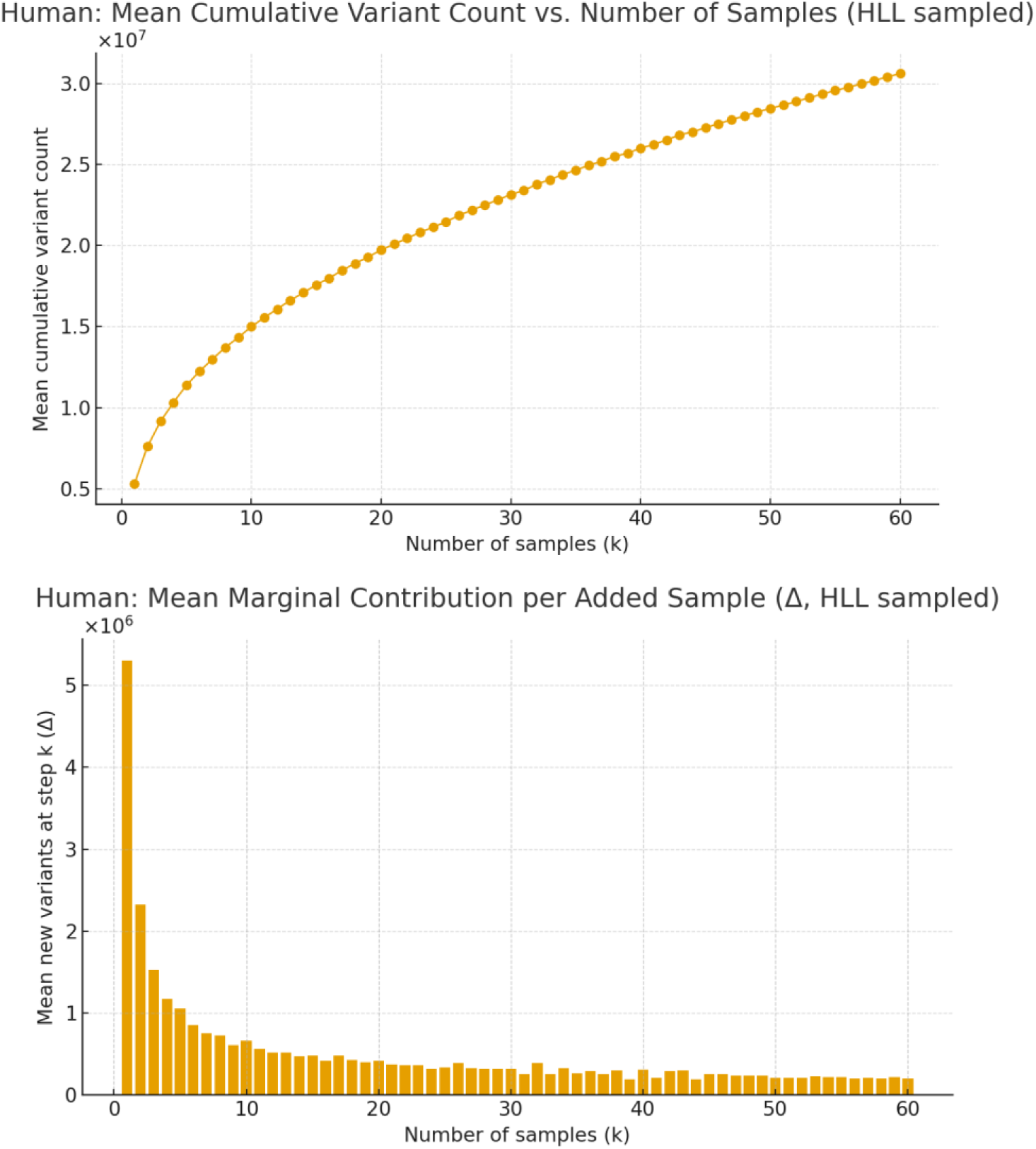
Cumulative unique variant counts as a function of the number of human genomes combined. **(a)** Mean cumulative variant count vs number of human genomes (out of 60) using HyperLogLog to estimate unique sites. The curve demonstrates how the total number of distinct variant loci grows with each additional genome. Starting at 5 million for one genome (the individual’s variants), the cumulative count reaches 30 million with 60 genomes and is still climbing, with no clear saturation at this cohort size. This indicates that many new variant sites are discovered as we sequence more individuals, reflecting the vast genetic diversity in humans. **(b)** Mean marginal new unique variants contributed per genome added. This shows the decrease in novel variants with each successive genome: the first genome adds 5×10^6^ unique sites, the second adds 2–3×10^6^ new ones, and by the 60th genome, each adds on the order of 10^4^–10^5^ new variant sites. The curve’s downward trend illustrates diminishing returns, yet a non-zero contribution even at genome 60; meaning every new individual, especially from a diverse background, still introduces some unique variants. Collectively, panels (a) and (b) highlight that large sample sizes are required to capture human genetic variation comprehensively; even 60 genomes are insufficient to reach a plateau. The data justify pangenomic reference models that incorporate many genomes, and they showcase the GPU pipeline’s capability to handle such multi-genome analyses.

Extrapolating from this trend suggests that substantially larger cohorts, potentially orders of magnitude larger than those analyzed here, would be required for the curve to approach a plateau, although formal statistical modelling would be required for precise projection. These findings demonstrate that scaling up to population-level genome cohorts is essential to capture the full spectrum of human genetic variation and that our GPU-accelerated pipeline makes it feasible to do so by processing many genomes rapidly. The results also reaffirm the inadequacy of any single reference genome for representing human diversity, since potentially tens of millions of loci lie outside one individual’s genome alignment to a singular reference and only become evident through a multi-genome (pangenome) analysis (Nyaga et al., 2025).

Among existing human datasets, the Simons Genome Diversity Project (SGDP; <u>Mallick et al., 2016</u>) most closely approximates an ecotype-style sampling strategy by prioritizing phylogenetic breadth over population depth. However, even SGDP cannot replicate the discrete lineage structure achievable in model organisms such as *C. elegans*, and most SGDP samples are distributed predominantly in derived formats (BAMs and VCFs) rather than uniformly accessible raw FASTQ files, constraining full recomputation under evolving reference models and algorithms.

### Whole Genome Variant Density Patterns

The distribution of variants across the human genome was investigated by binning variant calls into 10 Mb intervals and comparing cumulative variant densities for different cohort sizes (Fig. 5). In Fig. 5a, using a subset of 5 human genomes, we observe that variant density is already non-uniform across chromosomes. Certain regions harbour higher concentrations of variants, while others have relatively few. These peaks likely correspond to genomic regions known for high diversity or complex genomic content (for instance, the MHC region on chromosome 6, subtelomeric regions, or large gene families) (Logsdon et al., 2025), whereas troughs might represent more conserved or low-complexity regions. As the sample size increases to 20 genomes (Fig. 5b) and 50 genomes (Fig. 5c), the overall variant counts in each bin rise (note the increasing y-axis scales), but the general pattern of where the peaks and troughs are remains consistent. In other words, adding more genomes tends to amplify the variant counts across all regions rather than creating entirely new hotspot regions. By 50 genomes, virtually every 10 Mb window in the genome contains a substantial number of variants (on the order of tens of thousands), reflecting the ubiquity of genetic variation. However, the highest-density areas (peaks reaching >150,000 variants per 10 Mb) stand out even more with 50 genomes, indicating that these regions accumulate a disproportionate share of novel variants as cohorts grow. This suggests that some loci are highly polymorphic in the human population and continue to yield new variants with each added genome, whereas other regions become saturated quickly. The stable landscape of variant density from 5 to 50 genomes also implies that our variant calling is consistent and not biased by sample size. Regions that are variant-poor remain so even as more samples are added (indicating no artifactual calls under increased data). Together, Fig. 5a-c illustrate how large-scale genome sequencing reveals known diversity patterns: variant richness varies by genomic location and increases uniformly with sample size, requiring large cohorts to fully catalog the most variable regions of the genome.

**Figure 5:**
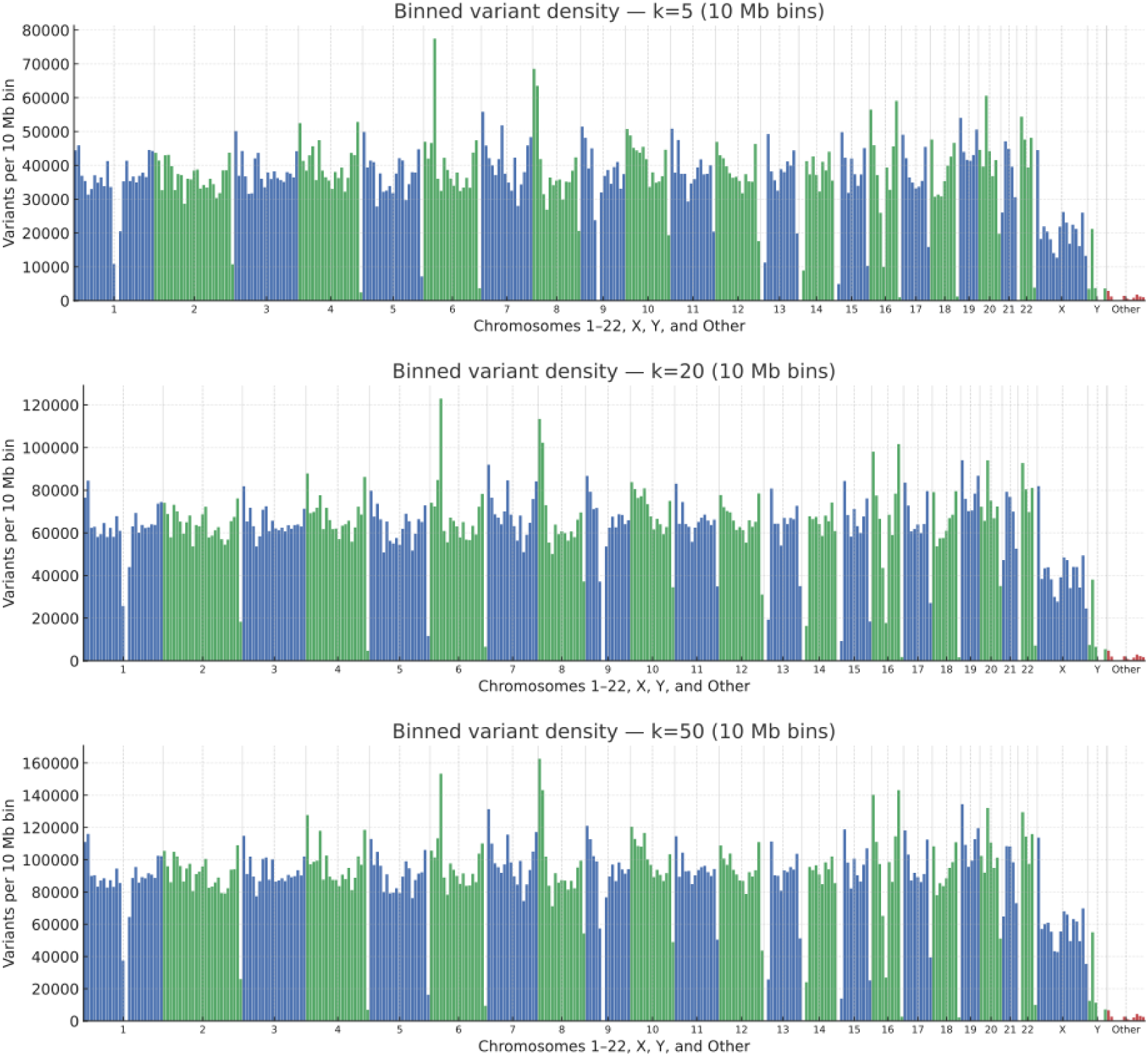
Genome-wide distribution of variant density for combined human cohorts of increasing size. Variant calls from multiple genomes were merged and binned into 10 Mb windows along each chromosome to visualize how variant density patterns change as more genomes are added. Three cohort sizes are shown: **(a)** 5 genomes, **(b)** 20 genomes, and **(c)** 50 genomes. The x-axis represents genomic position in order (chromosomes 1–22, X, Y, and “Other” unplaced contigs), and the y-axis indicates the number of variant sites in each 10 Mb bin (note the differing scales in a, b, c). At 5 genomes (a): key variant-rich regions are already evident as peaks (for instance, on chr6 likely corresponding to the hyper-variable HLA region, and subtelomeric regions on various chromosomes), whereas gene-poor or highly conserved regions show lower variant counts. At 20 genomes (b): overall variant counts roughly double compared to 5 genomes, and every chromosome accrues more variants, but the relative pattern of peaks and valleys remains similar – high-diversity regions accumulate variants from multiple individuals and still stand out. At 50 genomes (c): variant counts roughly scale with the number of genomes (many bins now contain well over 100,000 variants), and virtually no 10 Mb interval is devoid of variation. The highest peaks (e.g., in panel c some bins exceed 150k variants) correspond to loci that continue to gain new rare variants even as sample size grows. In contrast, some consistently low-variant regions (valleys) suggest areas of the genome that are universally conserved or less prone to common variation. Overall, Figure 5 shows that adding more genomes increases variant density across the board, but the shape of the variant landscape is intrinsic to the genome – certain hotspots of genetic variation are highlighted when enough genomes are surveyed. This also validates that the variant calling pipeline performs uniformly, as no new artifacts appear with larger cohorts; regions that were variant-sparse with 5 genomes remain so with 50, indicating reliable calls rather than noise.

### Population Ancestry and Variant Yield

To investigate human genetic diversity in the context of ancestry, we grouped the 60 genomes by continental heritage and compared their variant call yields (Table 5). All groups had comparable sequencing depth (37×–38× coverage on average) and similar runtime (35–37 minutes per genome), indicating uniform pipeline efficiency regardless of ancestry. However, there were clear differences in the average number of variants per genome among populations (Table 5). African-ancestry individuals (AFR, n=15) showed the highest variant counts, averaging approximately 5.53 million variants per genome. In contrast, East Asian genomes (EAS, n=13) had the lowest diversity with around 4.79 million variants on average, and Europeans (EUR, n=10) were slightly higher (4.89 million). South Asians (SAS, n=14) and Admixed Americans (AMR, n=8) fell in between (5.29M and 5.12M, respectively). These results are in line with established human population genetics: African genomes are known to harbor greater genetic variation than those from other continents, owing to Africa being the cradle of human genetic diversity (Sibomana, 2024). Indeed, the observed gap of 0.7 million variants between the average African and East Asian genome here is consistent with reports that African individuals carry 15–20% more variants than European or Asian individuals on average (Sibomana, 2024). The fact that our pipeline detects these differences underscores its sensitivity and accuracy in capturing true biological variation. Importantly, the GPU acceleration enables such multi-sample analyses to be done conveniently. For instance, an entire 60-genome panel was processed in under 36 hours (60 genomes × 0.6 h each), whereas on CPU the same would require 906 hours cumulatively, making this kind of comparative analysis far more labor-intensive without acceleration. Table 5 overall highlights that by coupling fast WGS analysis with diverse cohort sampling, we can readily characterize population-specific genomic diversity, which has implications for equitable genomic medicine and variant discovery in underrepresented groups.

**Table 5:** Performance and variant call summary by continental ancestry group (human genomes). This table stratifies the 60 human WGS samples into five heritage groups (African (AFR), Admixed American (AMR), East Asian (EAS), European (EUR), South Asian (SAS)) and shows the average coverage, read count, GPU runtime, and variant count per genome for each group. Coverage and reads are 38× and 750 million for all groups, and correspondingly, the average GPU time is 35–37 minutes per sample across ancestries; indicating no significant pipeline performance differences by sample ethnicity. In contrast, the average number of variants per genome varies by ancestry in line with known genetic diversity trends. African genomes average 5.53×10^6^ variants each, the highest among groups, consistent with Africa’s greater genetic diversity. South Asian and Admixed-American (AMR) genomes also show high variant counts (5.29M and 5.12M, respectively). European genomes average 4.89M, and East Asian genomes are lowest at 4.79M variants on average. These differences (on the order of 0.5–0.7 million variants between groups) illustrate the impact of demographic history on genetic variation captured per genome. Table 5 confirms that the GPU pipeline reliably captures these subtleties. It also emphasizes the importance of analyzing genomes from diverse populations: relying on a single reference or population could miss variants, whereas a multi-population approach uncovers a broader spectrum of variants. The overall average for all 60 samples (5.15M variants) is provided for reference. All groups had similar compute times and read counts, underscoring that the variant count differences are biological, not technical, and the accelerated pipeline is uniformly applicable to any population.

| Heritage<br>(Continent) | Number<br>of<br>Samples | Avg.<br>Coverage | Avg. Reads<br>Processed | Avg.<br>Compute<br>Time<br>(min)<sup>1</sup> | Avg.<br>Variants<br>Called |
| --- | --- | --- | --- | --- | --- |
| **African (AFR)** | 15 | 37.3x | 746 M | 34.8 min | 5,532,000 |
| **Admixed**<br>**American (AMR)** | 8 | 38.3x | 765 M | 36.2 min | 5,124,000 |
| **East Asian (EAS)** | 13 | 37.6x | 752 M | 34.9 min | 4,785,000 |
| **European (EUR)** | 10 | 38.5x | 769 M | 35.6 min | 4,888,000 |
| **South Asian (SAS)** | 14 | 37.9x | 758 M | 36.7 min | 5,286,000 |
| **Overall Average** | **60** | **37.7x** | **756 M** | **35.8 min** | **5,145,000** |

### Cost and Throughput Efficiency

Finally, we evaluated the computational cost implications of GPU acceleration in a cloud computing scenario (Table 6). Despite the higher hourly price of a GPU instance, the vast speedup yields a net cost advantage for the GPU pipeline. Using current AWS pricing, a 96-vCPU cloud instance (m6i.24xlarge) costs around $4.60 per hour, whereas a multi-GPU instance (g5.48xlarge, containing 8 NVIDIA A10 GPUs) is about $16.30 per hour on-demand. Processing a single human genome on the CPU instance took 15.1 hours, incurring an estimated cost of $17.37 per sample. In contrast, the GPU instance processed a genome in 0.59 hours (35 minutes), costing only $9.62 in that time (Table 6). This equates to roughly 45% lower cost per genome with the GPU pipeline on on-demand pricing, in addition to the 26× time savings. The cost efficiency gap widens even further if one leverages *spot instances* (preemptible transient cloud instances) for the GPU, which can be priced as low as 10% of on-demand rates. Using prices/discounts at the time of writing ($1.63/hour for the g5.48xlarge used in this paper), the cost per genome comes down to a remarkable <$1 per sample ($0.96). In this optimal scenario, the GPU pipeline is 18× cheaper than the CPU pipeline, utterly flipping the conventional wisdom that faster GPU computing is always more expensive. Even acknowledging the potential variability of spot instance availability, the ability to complete a genome analysis in well under an hour means one can opportunistically use short-lived cheap instances with high success. These findings demonstrate that GPU-accelerated genomics can be highly cost-effective, especially for embarrassingly parallel workloads. By reducing both runtime and cost per sample, the approach makes it feasible to scale WGS analysis to thousands of genomes. Moreover, the minimized intermediate data and shorter compute times also imply lower storage and energy footprints per genome analyzed. Overall, Table 6 underlines that investing in accelerated computing can yield substantial economic benefits for large-scale genomics projects, not only speeding up discoveries but doing so at lower cost than legacy CPU-based pipelines (O’Connell et al., 2023).

**Table 6:** Compute time and cost comparison of CPU vs GPU pipelines for human WGS (cloud instance example). Table contrasting the performance and cost per genome between a 24×large CPU-only AWS instance and a 48×large GPU-enabled AWS instance, including an analysis of using spot pricing for the GPU. Avg. Time/Sample: On the CPU instance (m6i.24xlarge, 96 vCPUs) the pipeline will take 15.1 hours per genome, whereas on the GPU instance (g5.48xlarge, 8× NVIDIA A10 GPUs) it took 0.59 hours (35 minutes). Approx. Hourly Cost: Lists the typical on-demand hourly rate for each instance type (CPU $4.60/hr, GPU $16.30/hr, and GPU spot $1.63/hr, which is 10% of on-demand). Approx. Cost/Sample: CPU cost/sample is computed as (hours × hourly cost) / 4 because 4 genomes run concurrently; GPU costs are hours × hourly cost. The CPU pipeline cost about $17.37 per genome, the GPU on-demand about $9.62 per genome, and the GPU on a spot instance only $0.96 per genome. Thus, even at on-demand pricing the GPU pipeline is nearly half the cost of the CPU pipeline for a given genome, thanks to the enormous time reduction. Using discounted spot instances, the cost drops by another order of magnitude, making high-throughput genome analysis extremely economical (less than $1 in compute spend per genome). These findings illustrate that GPU acceleration can simultaneously achieve faster turnaround and lower costs. The cost benefits are most pronounced when GPUs are used opportunistically (spot instances or otherwise high-utilisation scenarios). In practical terms, this means an entire 30× human genome can be analyzed for under a dollar of cloud compute, which has transformative implications for population-scale genomics (enabling thousands of genomes to be processed within typical research budgets). Footnotes: ¹m6i.24xlarge: Intel Xeon-based 96 vCPU instance (baseline for CPU pipeline). ²g5.48xlarge: Instance with 8× NVIDIA A10 Tensor Core GPUs (used for GPU runs). ³Spot pricing assumes 10% of on-demand cost (actual spot prices may vary).

| **Metric** | **CPU**<br>**(m6i.24xlarge)** | **GPU (g5.48xlarge)** | **GPU Spot**<br>**(g5.48xlarge)** |
| --- | --- | --- | --- |
| **Avg. Compute**<br>**Time/Sample** | 15.1 hours | 0.59 hours (35 min) | 0.59 hours (35 min) |
| **Approx. Hourly**<br>**Cost** | \$4.60 | \$16.30 | \$1.63 (10% of OD) |
| **Approx.**<br>**Cost/Sample** | **\$17.37** | **\$9.62** | **\$0.96** |

## Discussion

Embarrassingly_FASTA demonstrates that whole-genome preprocessing can be made both fast and economically scalable without obvious differences in variant-yield metrics (counts),. Across eight 30× human whole-genome samples, GPU execution reduced average end-to-end runtime from 15.1 hours using a CPU workflow with 24 threads per sample with 4 concurrent samples on a 96-vCPU node to 0.58 hours (36 minutes), delivering 26× higher throughput while producing nearly identical variant counts (5.1 million variants per genome) with differences of <0.3% and no systematic bias. This performance generalized across organisms: *C. elegans* genomes were processed in minutes (mean 4.7 min; <10 min worst case) and human genomes in 36 minutes on average, with runtimes scaling sub-linearly with read depth and remaining stable across globally distributed ecotypes and human heritages. These results confirm that GPU-accelerated preprocessing can preserve analytical integrity while radically improving throughput.

Crucially, the operational consequence of sub-hour genome processing is not merely faster pipelines, but a fundamental change in how genomic data can be managed. When aligned and derived intermediates (e.g., BAM and VCF files) can be regenerated on demand in minutes rather than days, they no longer need to be treated as archival dependencies. Instead, raw FASTQ data can be retained as the canonical source, mitigating irreversible information loss and reference dependence that otherwise constrain reanalysis as reference genomes, including pangenomes, evolve (Liao et al., 2023; Nyaga et al., 2025). This recomputable paradigm enables repeated, systematic reinterpretation of the same genomic data under improved aligners, variant callers, and reference models (Baykal et al., 2024) without prohibitive cost or delay. Our goal is not to outperform proprietary accelerators such as DRAGEN or Sentieon (Kendig et al., 2019; Samarakoon et al., 2025) at the kernel level, but to demonstrate a system architecture that fundamentally changes the economics, recomputability, and scalability of genome processing in cloud environments.

This shift in turn enables a step change in cloud economics for genomics. Using current AWS pricing, GPU processing reduced per-genome cost even under on-demand rates, and spot pricing lowered compute-only cost to below $1 per genome in our configuration, converting what was historically a multi-day, high-cost workflow into a short, preemption-tolerant job well suited to ephemeral infrastructure. With this capability, we performed a simulated pangenome build-up to quantify how variant diversity accumulates as non-redundant genomes are added. In *C. elegans*, cumulative variant discovery showed strong diminishing returns by 100 ecotypes (3.5-3.7 million unique variant sites), with marginal gains dropping to only a few thousand new sites per additional genome. In humans, 60 genomes spanning five continental heritage buckets yielded ∼30 million unique variant sites with no clear plateau at this cohort size, and ancestry-stratified results recapitulated known diversity patterns, including elevated per-genome variant counts in African-ancestry samples (1000 Genomes Project Consortium et al., 2015; Mallick et al., 2016; Sibomana, 2024). Notably, if *C. elegans* ecotypes were artificially collapsed into a small number of broad geographic groupings analogous to human continental buckets, its diversity curve would likewise show weaker flattening over the same number of samples, indicating that the observed contrast reflects sampling granularity rather than intrinsic limits on species-level diversity. Together, these findings underscore that human genetic diversity remains far from exhaustively sampled under current cohort sizes and coarse heritage bucketing, and that progress toward comprehensive catalogs requires both larger cohorts and the ability to reprocess them repeatedly as references and algorithms improve (Liao et al., 2023; Mallick et al., 2016).

Beyond immediate performance and cost gains, Embarrassingly_FASTA directly addresses a central barrier to the emergence of AI-enabled World Genome Models (WGMs): the ability to repeatedly and economically recompute population-scale genomic representations as references, algorithms, and cohort composition evolve. WGMs require not only vast volumes of genomic data, but a computational substrate that preserves access to raw sequence while supporting continual reprocessing across linear references, pangenomes, and future representations. By rendering intermediate files transient and enabling rapid, low-cost recomputation from FASTQ, our approach shifts genomic data management from a static, archival paradigm to a dynamic, recomputable one. This is essential for training and validating WGMs, which must integrate millions of genomes across ancestries while remaining adaptable to new reference models, improved alignment strategies, sparse and dense genomic representations, and evolving notions of genomic “ground truth.” In this sense, Embarrassingly_FASTA is not merely a faster pipeline, but enabling infrastructure for WGM-scale genomics, making it practical to curate, revise, and expand the genomic corpora on which next-generation disease-mechanism discovery and population-scale genomic AI will depend.

### Limitations

While *Embarrassingly_FASTA* demonstrates a scalable path toward recomputable genomics, our study has several limitations. First, our cost efficiency analysis relies on cloud spot instances, whose availability and pricing are market-driven and can fluctuate regionally. We do not implement explicit checkpointing or interruption-aware orchestration; rather, the pipeline is preemption-tolerant primarily because per-sample runtimes are sub-hour, making simple retries economically viable. Second, the current implementation aligns reads to a linear reference genome (GRCh38) and does not yet incorporate the additional computational overhead of aligning directly to pangenome graphs. Third, the CPU and GPU workflows used different variant calling algorithms (BCFtools vs. GATK HaplotypeCaller via Parabricks); the small differences observed likely reflect algorithmic differences rather than hardware effects, and rigorous correctness evaluation would require truth-set benchmarking or site-level concordance.

## Materials and Methods

### Data Sources and Reference Genomes

We benchmarked both pipelines using real whole-genome sequencing (WGS) data from two species. For *Caenorhabditis elegans*, we obtained 100 WGS samples from the Caenorhabditis Natural Diversity Resource (CaeNDR) database (Crombie et al., 2024). For human data, we used 60 WGS samples from the 1000 Genomes Project (1000 Genomes Project Consortium et al., 2015). All raw read data (FASTQ files) were downloaded from the European Nucleotide Archive (ENA) using the accession information provided by those projects. All analyses comply with the original consent and data use policies of the respective consortia.

We used the latest reference genomes appropriate for each species. Human reads were aligned to the Ensembl GRCh38 reference genome (Release 104), and *C. elegans* reads were aligned to the WBcel235 reference genome (Release 115). Both reference genome builds were obtained from Ensembl/WormBase FTP repositories, including all associated index files. Prior to alignment, each reference genome was indexed using the appropriate aligner (BWA or Parabricks) index commands.

### Cost Analysis and Commercial Benchmarking

To establish a baseline for current commercial genomic processing costs, we conducted a market survey of independent bioinformatic service providers in January 2026. We contacted seven service providers to request quotes for the secondary analysis (alignment and variant calling) of 60 human WGS samples (30X coverage), representative of the workload presented in this study. Of the seven providers contacted, three did not reply. The remaining four providers returned quotes with an average processing cost of $121 per sample. This commercial baseline was compared against our internal cloud costs calculated using Amazon Web Services (AWS) pricing for the specific instance types utilized (m6i.24xlarge for CPU and g5.48xlarge for GPU). Cloud costs were derived from both standard on-demand rates and spot instance pricing active at the time of the experiments. AWS instance prices were calculated using the US East (N. Virginia) region spot and on-demand pricing as of January 2026.

### CPU-Based Variant Calling Pipeline

All steps of the CPU pipeline were implemented in Python 3.9 and orchestrated through a Bash wrapper script (run_pipeline.sh) calling a Python workflow script (pipeline_script.py). The pipeline processed each sample’s reads through alignment, post-processing, and variant calling as described below:

#### Read Alignment

Paired-end reads were aligned to the reference genome using BWA-MEM (Li & Durbin, 2009). BWA-MEM (v0.7.17) was run with default parameters to produce SAM alignments for each sample.

#### Sorting and Indexing

The raw SAM alignments were converted to BAM format, then sorted and indexed using SAMtools (v1.23) (Danecek et al., 2021; Li et al., 2009). This ensured reads were ordered by coordinate and allowed rapid random access for downstream steps.

#### Duplicate Marking

We marked PCR duplicates in each sorted BAM using Picard MarkDuplicates (*Picard Tools - By Broad Institute*, n.d.). Picard (v3.4.0) also generated alignment metrics (e.g. percentage duplicates) as quality control. The resulting BAM files had duplicate reads flagged but not removed.

#### Variant Calling

Variants were called from each de-duplicated BAM using **BCFtools (**v1.23) in two steps: bcftools mpileup to generate genotype likelihoods, and bcftools call to call SNPs and indels. This calling approach implements the original Samtools/BCFtools variant calling method (Li, 2011). Each sample’s variants were output to a VCF file.

#### Resource Monitoring

Throughout the CPU pipeline, resource usage was tracked. We used the psutil Python library to record CPU and memory utilization at each step, and the Linux iostat tool to monitor disk I/O throughput. This allowed profiling of the CPU pipeline’s performance characteristics on the given hardware.

#### GPU-Accelerated Variant Calling Pipeline

For GPU-accelerated processing, we utilized NVIDIA Clara Parabricks (Zhu et al., 2025) version 4.5.1-1, a suite of GPU-optimized genome analysis tools provided by NVIDIA. The GPU pipeline was orchestrated with Python 3.12 scripts (gpu_pipeline_AWS.py with helper functions in pipeline_functions_AWS.py), which launched Parabricks container jobs on the AWS GPU instance. The workflow for each sample was:

#### FASTQ to Aligned BAM

We used the Parabricks pbrun fq2bam command to perform read alignment and post-processing in one step. This tool aligns reads to the reference genome (using an NVIDIA-optimized equivalent of BWA-MEM), sorts the alignments, marks duplicates, and produces a sorted, indexed BAM. The pbrun fq2bam stage is functionally analogous to steps 1–3 of the CPU pipeline but leverages GPU acceleration inside a Docker container.

#### Variant Calling

We called variants using Parabricks’ GPU-accelerated implementation of the GATK HaplotypeCaller (v4.6.1.0). Specifically, the pipeline invoked pbrun haplotypecaller to generate per-sample VCF files of variant calls from each BAM. This tool parallels GATK HaplotypeCaller logic but is optimized for NVIDIA GPUs within the Parabricks environment.

#### Pipeline comparability and algorithmic considerations

Although the CPU and GPU pipelines employ different variant callers (BCFtools on CPU and GATK HaplotypeCaller via Parabricks on GPU), the near-identical variant yields and lack of systematic bias observed between pipelines indicate that performance improvements are dominated by hardware acceleration rather than algorithmic divergence. This is further supported by prior benchmarking studies demonstrating high concordance between modern variant callers across diverse datasets, and by the fact that both pipelines implement probabilistic SNP and indel calling under comparable assumptions (Supernat et al., 2018). We selected BCFtools for the CPU baseline specifically because it is demonstrated to be significantly faster than CPU-based GATK HaplotypeCaller (Lefouili & Nam, 2022). Consequently, our reported 26× speedup represents a conservative lower bound; a direct comparison to the computationally heavier CPU-GATK would have resulted in an even larger performance gap.

#### GPU Resource Monitoring

During the GPU pipeline runs, we collected GPU utilization and memory metrics using nvidia-smi at regular intervals. These metrics captured GPU memory consumption and compute utilization for fq2bam and haplotypecaller processes, enabling a performance comparison with the CPU pipeline.

All GPU pipeline steps were executed inside NVIDIA Parabricks Docker containers. The Python orchestration script managed container execution and data movement (mounting input FASTQ files and saving output BAM/VCF files to shared storage). This ensured a reproducible GPU workflow encapsulated in containers.

### System-Level Contributions of Embarrassingly_FASTA Beyond Parabricks

Embarrassingly_FASTA is not a new variant calling algorithm, but a systems-level framework/deployment that re-architects GPU-accelerated genomics around transient, fault-tolerant, and economically scalable execution. While Parabricks provides GPU-optimized kernels for alignment and variant calling, Embarrassingly_FASTA contributes:

i. **GPU-pinned concurrent execution** enabling cohort-level packing (e.g., running multiple small-genome samples concurrently via fixed GPU partitioning).
ii. **Tunable + instrumented Parabricks deployment** (e.g., gpuwrite/gpusort/nstreams/queue tuning) with telemetry for CPU/GPU/I/O bottleneck attribution.
iii. **Transient-intermediate lifecycle** that treats BAM/VCF as regenerable artifacts and retains FASTQ as canonical input for recomputation.

These capabilities enable a class of genomic workflows that are impractical under traditional persistent cluster or on-demand cloud assumptions.

### Computational Resources

We ran CPU and GPU pipelines on Amazon Web Services (AWS) EC2 instances to ensure hardware environment consistency. For the GPU pipeline, we used the g5.48xlarge instance type, which provides 8 x NVIDIA A10 Tensor Core GPUs along with 192 vCPUs and 768 GB of RAM (AWS g5 family). The CPU pipeline was run on a high-end CPU instance, m6i.24xlarge, which provides 96 vCPUs (Intel Xeon Ice Lake processors) and 384 GB of RAM, but no GPU. Both instance types were running the Linux (Version AL2023) operating system and accessed the same input data from attached high-throughput storage. Only 2 GPUs were utilised for the *C. elegans* processing, whereas the full 8 GPUs were used for *H. sapiens.* In data not shown, using more than 2 GPUs for *C. elegans* processing did not result in any performance improvement, thus highlighting a maximum speed gain in parallelisation.

Each pipeline was executed on its respective instance type for all samples to measure wall-clock time and resource utilization. We repeated runs as needed to account for variability and ensured no other major workloads were running on the instances during benchmarking.

### Downstream Variant Analysis

After obtaining variant call sets for all samples, we conducted additional analyses to compare results and evaluate genomic coverage of variants:

#### Variant Set Extraction

For each sample’s VCF, we extracted the set of variant positions using **bcftools query**. This yielded a list of unique variants (by genomic coordinate and allele) per sample. These per-sample variant lists served as the basis for diversity and union analysis.

#### Cumulative Union of Variants

Using a custom Python script (cumulative_union.py), we iteratively combined variant sets across samples to assess the accumulation of unique variants. In other words, we started with one sample’s variant set and progressively took the union with each additional sample’s variants, tracking the total number of unique variants after each union step. This analysis was done separately for the CPU pipeline VCFs and the GPU pipeline VCFs to compare how each pipeline captured genetic diversity as more genomes were added.

#### Diversity Estimation with HyperLogLog

To efficiently estimate the cardinality of unions for large numbers of samples, we implemented a HyperLogLog (HLL) probabilistic counting approach. We used the HyperLogLog algorithm(Heule et al., 2013) via the datasketch (v1.6.5) Python library (which internally uses the xxHash hashing algorithm for speed). A script (mean_k_unions_hll.py) was used to estimate the expected number of unique variants when considering combinations of *k* samples, without having to explicitly merge all variants for every subset. This HLL-based approach gave an approximate but scalable measure of variant diversity across the cohort, and we validated it against exact counts for smaller *k* values.

#### Genomic Distribution of Variants

We examined how variants were distributed across the reference genome by binning variant calls into 10 Mb windows. Using a custom visualization script (sandbox_density_vs_searchspace.py), we plotted variant density (number of variants) versus genomic position for each pipeline’s calls. This helped identify any coverage gaps or biases in variant calls along each chromosome. The script took the variant coordinate lists and computed densities per 10 Mb bin, producing comparative plots for CPU vs. GPU pipeline results.

All analysis scripts were written in Python 3.12 and took advantage of libraries like pandas (The pandas development team, 2025) for data manipulation and numpy (*Array Programming with NumPy | Nature*, n.d.) for numerical calculations. The results of these downstream analyses were used to quantify differences in variant calling completeness and consistency between the two pipelines.

### Statistical Analysis and Visualization

We created publication-quality figures using Python-based tools. Timing and resource usage logs from both pipelines were parsed with pandas (v3.0.0), and summary statistics (mean, median, etc.) were computed to compare CPU vs. GPU performance. All plots were generated with Matplotlib (Hunter, 2007) (v3.7) and styled for clarity. Bar charts were used to compare average runtimes and resource utilization between the CPU and GPU pipelines. Line plots were used for the cumulative variant union analysis, showing how the count of unique variants grows as more genomes are added. We also generated world map visualizations to show the geographic origin of the samples: using our map_plot.py script, we leveraged the Cartopy (v0.25.0) (Elson et al., 2024) library with Natural Earth datasets to draw world maps, marking the sampling locations of the *C. elegans* strains and human populations included in the study. This provided context on the diversity of samples and ensured that our dataset covered multiple continents. All figure scripts (including map plotting and density plots) are available in the project repository.

## Code Availability and Reproducibility

All pipeline scripts, analysis scripts, and Docker container configurations used in this study are available in our project’s GitHub repository https://github.com/ecotoneservice/Embarrassingly_FASTA (including pipeline_script.py, gpu_pipeline_AWS.py, and supporting modules). The repository contains a README with instructions for reproducing the CPU and GPU pipeline runs, including environment setup and required dependencies (install_dependencies.py). By providing the exact versions of tools (BWA-MEM, SAMtools, Picard, BCFtools, Parabricks, etc.) and the analysis scripts used, we ensure that the benchmarking results are reproducible. The Docker images for the GPU pipeline (NVIDIA Clara Parabricks v4.5.1-1) can be obtained via NVIDIA’s container registry as documented in our repository. All data processing steps described above can be re-run by interested researchers to verify the findings or adapt the pipelines to their own data.

## Notes

https://github.com/ecotoneservice/Embarrassingly_FASTA

### Competing Interest Statement

The authors have declared no competing interest.

### Summary of Updates

Minor formatting and typographical changes between v1 and v2.

## References

1000 Genomes Project Consortium, Auton, A., Brooks, L. D., Durbin, R. M., Garrison, E. P., Kang, H. M., Korbel, J. O., Marchini, J. L., McCarthy, S., McVean, G. A., & Abecasis, G. R. (2015). A global reference for human genetic variation. Nature, 526(7571), 68–74. 10.1038/nature15393

Array programming with NumPy | Nature. (n.d.). Retrieved November 20, 2025, from https://www.nature.com/articles/s41586-020-2649-2

Avsec, Ž., Latysheva, N., Cheng, J., Novati, G., Taylor, K. R., Ward, T., Bycroft, C., Nicolaisen, L., Arvaniti, E., Pan, J., Thomas, R., Dutordoir, V., Perino, M., De, S., Karollus, A., Gayoso, A., Sargeant, T., Mottram, A., Wong, L. H., … Kohli, P. (2025). AlphaGenome: Advancing regulatory variant effect prediction with a unified DNA sequence model (p. 2025.06.25.661532). bioRxiv. 10.1101/2025.06.25.661532

Baykal, P. I., Łabaj, P. P., Markowetz, F., Schriml, L. M., Stekhoven, D. J., Mangul, S., & Beerenwinkel, N. (2024). Genomic reproducibility in the bioinformatics era. Genome Biology, 25(1), 213. 10.1186/s13059-024-03343-2

Brixi, G., Durrant, M. G., Ku, J., Poli, M., Brockman, G., Chang, D., Gonzalez, G. A., King, S. H., Li, D. B., Merchant, A. T., Naghipourfar, M., Nguyen, E., Ricci-Tam, C., Romero, D. W., Sun, G., Taghibakshi, A., Vorontsov, A., Yang, B., Deng, M., … Hie, B. L. (2025). Genome modeling and design across all domains of life with Evo 2 (p. 2025.02.18.638918). bioRxiv. 10.1101/2025.02.18.638918

Clavell-Revelles, P., Reese, F., Carbonell-Sala, S., Degalez, F., Arnan, C., Oliveros, W., Palumbo, E., Perteghella, T., Guigó, R., & Melé, M. (2025). Long-read transcriptomics of a diverse human cohort reveals ancestry bias in gene annotation. Nature Communications, 16(1), 10194. 10.1038/s41467-025-66096-x

Crombie, T. A., McKeown, R., Moya, N. D., Evans, K. S., Widmayer, S. J., LaGrassa, V., Roman, N., Tursunova, O., Zhang, G., Gibson, S. B., Buchanan, C. M., Roberto, N. M., Vieira, R., Tanny, R. E., & Andersen, E. C. (2024). CaeNDR, the Caenorhabditis Natural Diversity Resource. Nucleic Acids Research, 52(D1), D850–D858. 10.1093/nar/gkad887

Dalla-Torre, H., Gonzalez, L., Mendoza-Revilla, J., Lopez Carranza, N., Grzywaczewski, A. H., Oteri, F., Dallago, C., Trop, E., de Almeida, B. P., Sirelkhatim, H., Richard, G., Skwark, M., Beguir, K., Lopez, M., & Pierrot, T. (2025). Nucleotide Transformer: Building and evaluating robust foundation models for human genomics. Nature Methods, 22(2), 287–297. 10.1038/s41592-024-02523-z

Danecek, P., Bonfield, J. K., Liddle, J., Marshall, J., Ohan, V., Pollard, M. O., Whitwham, A., Keane, T., McCarthy, S. A., Davies, R. M., & Li, H. (2021). Twelve years of SAMtools and BCFtools. GigaScience, 10(2), giab008. 10.1093/gigascience/giab008

Elson, P., Andrade, E. S. de, Lucas, G., May, R., Hattersley, R., Campbell, E., Comer, R., Dawson, A., Little, B., Raynaud, S., scmc72, Snow, A. D., lgolston, Blay, B., Killick, P., lbdreyer, Peglar, P., Wilson, N., Andrew, … Kirkham, D. (2024). SciTools/cartopy: REL: v0.24.1 [Computer software]. Zenodo. 10.5281/zenodo.13905945

Feng, H., Wu, L., Zhao, B., Huff, C., Zhang, J., Wu, J., Lin, L., Wei, P., & Wu, C. (2025). Benchmarking DNA foundation models for genomic and genetic tasks. Nature Communications, 16, 10780. 10.1038/s41467-025-65823-8

Franke, K. R., & Crowgey, E. L. (2020). Accelerating next generation sequencing data analysis: An evaluation of optimized best practices for Genome Analysis Toolkit algorithms. Genomics & Informatics, 18(1), e10. 10.5808/GI.2020.18.1.e10

Guio, H., Sanchez, C., Borda, V., Jaramillo-Valverde, L., Caceres, O., Padilla, C., Trujillo, O., Poterico, J. A., Silva-Carvalho, C., Horton, M., Lanata, C. M., Carnevale, A., Romero-Hidalgo, S., Canizales-Quinteros, S., Acuña-Alonzo, V., Machacuay-Romero, M., Novoa, P., Frisancho, R., Shady, R., … Tarazona-Santos, E. (2025). The peruvian genome project: Expanding the global pool of genome diversity from South America. Frontiers in Genetics, 16, 1614021. 10.3389/fgene.2025.1614021

Heule, S., Nunkesser, M., & Hall, A. (2013). HyperLogLog in Practice: Algorithmic Engineering of a State of The Art Cardinality Estimation Algorithm. 683–692. http://stefanheule.com/publications/edbt13-hyperloglog/

Hunter, J. D. (2007). Matplotlib: A 2D Graphics Environment. Computing in Science & Engineering, 9(3), 90–95. 10.1109/MCSE.2007.55

*Illumina’s revolutionary NovaSeq X exceeds 200th order milestone in first quarter* 2023. (n.d.). Retrieved December 11, 2025, from https://www.illumina.com/content/illumina-marketing/amr/en_US/company/news-center/press-releases/2023/4b48580b-42d4-419b-8512-5adcbb069836.html?null

Jeon, S., Choi, Hansol, Jeon, Y., Choi, W.-H., Choi, Hyunjoo, An, K., Ryu, H., Bhak, Jihun, Lee, H., Kwon, Y., Ha, S., Kim, Y. J., Blazyte, A., Kim, C., Kim, Y., Kang, Y., Woo, Y. J., Lee, C., Seo, J., … Bhak, J. (2024). Korea4K: Whole genome sequences of 4,157 Koreans with 107 phenotypes derived from extensive health check-ups. GigaScience, 13, giae014. 10.1093/gigascience/giae014

Kendig, K. I., Baheti, S., Bockol, M. A., Drucker, T. M., Hart, S. N., Heldenbrand, J. R., Hernaez, M., Hudson, M. E., Kalmbach, M. T., Klee, E. W., Mattson, N. R., Ross, C. A., Taschuk, M., Wieben, E. D., Wiepert, M., Wildman, D. E., & Mainzer, L. S. (2019). Sentieon DNASeq Variant Calling Workflow Demonstrates Strong Computational Performance and Accuracy. Frontiers in Genetics, 10, 736. 10.3389/fgene.2019.00736

Koreniuk, O., & Njie, eMalick G. (2025). dnaSORA - A Unified Diffusion Transformer for DNA point clouds (p. 2025.01.27.633223). bioRxiv. 10.1101/2025.01.27.633223

Lefouili, M., & Nam, K. (2022). The evaluation of Bcftools mpileup and GATK HaplotypeCaller for variant calling in non-human species. Scientific Reports, 12(1), 11331. 10.1038/s41598-022-15563-2

Li, H. (2011). A statistical framework for SNP calling, mutation discovery, association mapping and population genetical parameter estimation from sequencing data. *Bioinformatics (Oxford*, England), 27(21), 2987–2993. 10.1093/bioinformatics/btr509

Li, H., & Durbin, R. (2009). Fast and accurate short read alignment with Burrows–Wheeler transform. Bioinformatics, 25(14), 1754–1760. 10.1093/bioinformatics/btp324

Li, H., Handsaker, B., Wysoker, A., Fennell, T., Ruan, J., Homer, N., Marth, G., Abecasis, G., Durbin, R., & 1000 Genome Project Data Processing Subgroup. (2009). The Sequence Alignment/Map format and SAMtools. Bioinformatics (Oxford, England), 25(16), 2078–2079. 10.1093/bioinformatics/btp352

Liao, W.-W., Asri, M., Ebler, J., Doerr, D., Haukness, M., Hickey, G., Lu, S., Lucas, J. K., Monlong, J., Abel, H. J., Buonaiuto, S., Chang, X. H., Cheng, H., Chu, J., Colonna, V., Eizenga, J. M., Feng, X., Fischer, C., Fulton, R. S., … Paten, B. (2023). A draft human pangenome reference. Nature, 617(7960), 312–324. 10.1038/s41586-023-05896-x

Lin, M.-J., Iyer, S., Chen, N.-C., & Langmead, B. (2024). Measuring, visualizing and diagnosing reference bias with biastools. bioRxiv: The Preprint Server for Biology, 2023.09.13.557552. 10.1101/2023.09.13.557552

Lloret-Villas, A., Bhati, M., Kadri, N. K., Fries, R., & Pausch, H. (2021). Investigating the impact of reference assembly choice on genomic analyses in a cattle breed. BMC Genomics, 22, 363. 10.1186/s12864-021-07554-w

Logsdon, G. A., Ebert, P., Audano, P. A., Loftus, M., Porubsky, D., Ebler, J., Yilmaz, F., Hallast, P., Prodanov, T., Yoo, D., Paisie, C. A., Harvey, W. T., Zhao, X., Martino, G. V., Henglin, M., Munson, K. M., Rabbani, K., Chin, C.-S., Gu, B., … Marschall, T. (2025). Complex genetic variation in nearly complete human genomes. Nature, 644(8076), 430–441. 10.1038/s41586-025-09140-6

Mallick, S., Li, H., Lipson, M., Mathieson, I., Gymrek, M., Racimo, F., Zhao, M., Chennagiri, N., Nordenfelt, S., Tandon, A., Skoglund, P., Lazaridis, I., Sankararaman, S., Fu, Q., Rohland, N., Renaud, G., Erlich, Y., Willems, T., Gallo, C., … Reich, D. (2016). The Simons Genome Diversity Project: 300 genomes from 142 diverse populations. Nature, 538(7624), 201–206. 10.1038/nature18964

Nyaga, D. M., Zaied, R. E., Silander, O. K., Black, M. A., & O’Sullivan, J. M. (2025). Beyond single references: Pangenome graphs and the future of genomic medicine. Frontiers in Genetics, 16. 10.3389/fgene.2025.1679660

O’Connell, K. A., Yosufzai, Z. B., Campbell, R. A., Lobb, C. J., Engelken, H. T., Gorrell, L. M., Carlson, T. B., Catana, J. J., Mikdadi, D., Bonazzi, V. R., & Klenk, J. A. (2023). Accelerating genomic workflows using NVIDIA Parabricks. BMC Bioinformatics, 24(1), 221. 10.1186/s12859-023-05292-2

Picard Tools—By Broad Institute. (n.d.). Retrieved November 20, 2025, from https://broadinstitute.github.io/picard/

Samarakoon, P. S., Fournous, G., Hansen, L. T., Wijesiri, A., Zhao, S., Alex A, R., Nandi, T. N., Madduri, R., Rowe, A. D., Thomassen, G., Hovig, E., & Razick, S. (2025). Benchmarking accelerated next-generation sequencing analysis pipelines. Bioinformatics Advances, 5(1), vbaf085. 10.1093/bioadv/vbaf085

Sibomana, O. (2024). Genetic Diversity Landscape in African Population: A Review of Implications for Personalized and Precision Medicine. Pharmacogenomics and Personalized Medicine, 17, 487–496. 10.2147/PGPM.S485452

Supernat, A., Vidarsson, O. V., Steen, V. M., & Stokowy, T. (2018). Comparison of three variant callers for human whole genome sequencing. Scientific Reports, 8, 17851. 10.1038/s41598-018-36177-7

Suzuki, Y. (2025). The First Advent of the Whole Genome Sequencing Cohort in Japan. JMA Journal, 8(4), 1053–1054. 10.31662/jmaj.2025-0389

The pandas development team. (2025). *pandas-dev/pandas: Pandas* [Computer software]. Zenodo. 10.5281/zenodo.17229934

Ultima Genomics | Ultima Genomics Delivers the $100 Genome. Emerges from Stealth with $600 Million in Funding. (n.d.). Retrieved December 11, 2025, from https://www.ultimagenomics.com/blog/ultima-genomics-delivers-the-usd100-genome/

Wetterstrand, K. (n.d.). Data from the NHGRI Genome Sequencing Program (GSP). Retrieved December 11, 2025, from https://www.genome.gov/about-genomics/fact-sheets/DNA-Sequencing-Costs-Data

Zhu, T., Vats, P., Onken, S., Dunstan, A., Zamirai, B., Puleri, D. F., Nair, A., Oliva, M., Gaihre, A., Sadhnani, P., Li, S., Arumugam, K., Chacon, A., Maric, M., Cohen, J., Sethia, A., & Samadi, M. (2025). *Parabricks: GPU Accelerated Universal Pan-Instrument Genomics Analysis Software Suite* (p. 2025.07.23.666378). bioRxiv. 10.1101/2025.07.23.666378

